# Quantum mechanics predicts Bicoid interpretation time of less than a second

**DOI:** 10.1101/2024.09.10.612267

**Authors:** Irfan Lone

**Affiliations:** Department of Physics and Astronomy, Howard University, Washington, DC 20059, USA; Canadian Quantum Research Center, 3002 32nd Ave Vernon, BC 204-3002 32, Canada; Department of Chemistry, Indian Institute of Technology Bombay, Powai, Mumbai 400076, India

## Abstract

The establishment and interpretation of the concentration gradient of the morphogen Bicoid (*Bcd*) is crucial for the successful embryonic development of fruit flies. However, the biophysical mechanisms behind the timely formation and subsequent interpretation of this prototypical morphogen gradient by its target genes are not yet completely understood. Recently a discrete time, one-dimensional quantum walk model of *Bcd* gradient formation has been successfully used to explain the observed multiple dynamic modes of the system. However, the question of its precise interpretation by its primary target gene hunchback (*hb*) remains still unanswered. In this paper it will be shown that the interpretation of the *Bcd* gradient by its primary target gene *hb*, with the observed precision of ∼ 10%, takes a time period of less than a second, as expected on the basis of recent experimental observations. Furthermore, the quantum walk model is also used to explain certain key observations of recent optogenetic experiments concerning the time windows for *Bcd* interpretation. Finally, it is concluded that the incorporation of quantum effects into the treatment of *Bcd* gradient represents a viable step in exploring its dynamics.

---

The establishment and subsequent interpretation of concentration gradients of certain signaling molecules, commonly called morphogens, plays a crucial role during the embryonic development of many multi-cellular organisms [1–5]. For instance, the concentration gradient of the transcription factor Bicoid (*Bcd*) established during the early embryonic development of fruit fly *Drosophila melanogaster* provides the necessary positional information and polarity for the proper implementation of the body plan of the organisms [6–10]. The timely establishment and precise interpretation of *Bcd* gradient by its primary target gene hunchback (*hb*) plays a crucial role in the successful development of the fruit fly.

Recent experiments, based on fluorescence correlation spectroscopy and perturbative techniques, have observed multiple dynamic modes accompanying the formation of the *Bcd* gradient [11]. These observations are best fitted by a one-dimensional quantum walk based model [12]. However, the biophysical mechanisms underlying the precise interpretation of *Bcd* by its primary target gene *hb*, with the observed precision of ∼ 10 %, still remain elusive [13–24]. Equilibrium models have essentially failed to capture this important aspect of the *Bcd*-*hb* system in a satisfactory manner and the formulation of increasingly more complex approaches has been suggested [17]. Theories employing detailed balance arguments have also not proven much helpful in this regard [25]. In recent years, therefore, theoretical treatments invoking the hypotheses of dynamically changing time windows have been carried out [26]. These predict an interpretation time of less than a minute and remain unsubstantiated in vivo [20, 21, 26]. However, experimentally *Bcd* protein is known to bind to chromatin sites in a highly transient manner, with specific binding events lasting on the order of a second, in all portions of the embryo [18]. These observations demand interpretation times of duration less than a second. Thus, despite extensive numerical and analytical work, the problem of interpretation of the *Bcd* morphogen by its primary target gene *hb* has no satisfactory solution.

The timescale of *Bcd* driven pattern formation in the fly embryo may arbitrarily be divided into two subintervals. First 90 minutes, prior to nuclear cycle 10, are utilized by the system for the establishment of the *Bcd* gradient in the embryo [27]. From nuclear cycle 10 onward the *Bcd* concentration inside the nuclei remains stable from cycle to cycle. It has been shown that the quantitative relationship between the statistics of *hb* expression levels in the embryo is consistent with a model in which the dominant source of noise are the diffusive fluctuations associated with the random arrival of *Bcd* transcription factor molecules at their target sites on the DNA [28]. This noise dependence is quantitatively expressed by the Berg-Purcell limit,

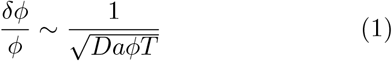

in which *a* denotes the size of the *Bcd* receptor, *T* is the duration of signal integration, *ϕ*(*x, t*) is the local *Bcd* concentration and *D* is the diffusion rate [29].

Intriguingly, what Gregor *et al*. discovered is that the observed precision of ∼ 10 % in the readout of the *Bcd* concentration by the nuclei is inconsistent with the predictions of this fundamental physical limit for diffusive processes [13]. To circumvent this problem, the idea of spatial averaging by the nuclei was introduced by the authors in order to increase the precision to the desired level. Over the years the mechanism of Gregor et al. has been widely applied [14–17] but the problem of interpretation of *Bcd* gradient by its primary target gene *hb* still remains poorly understood [17–24]. On the other hand, a recent theoretical analysis has shown that spatial averaging is a priori dispensable but the treatment introduces another *ad hoc* hypothesis of dynamically changing time windows [26]. Most importantly, it predicts an interpretation time much higher than is supported by current experimental observations based on lattice light sheet microscopy studies [18].

In this Letter we employ the results obtained through the quantum walk based analysis to predict the expected interpretation time of the *Bcd* morphogen by its primary target gene *hb* [13] and certain key observations of recent optogenetic experiments concerning the time windows for *Bcd* interpretation are also explained.

From the quantum walk based analysis one obtains the following expression for the local diffusivity of the *Bcd* gradient

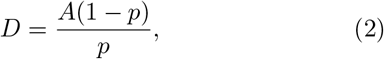

where *p* is a parameter that quantifies noise in the system and the coefficient *A* is a function of *p* with an estimated value of ∼0.40 and *A*→ 0.5 as *p*→ 1 [12]. Remarkably, Francis Crick, using his source-sink model,

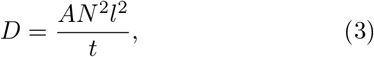

has estimated a value of 0.42 for *A* and suggested a general value of *A* = 0.5 for the biochemically more realistic models; where *t* denotes the time, *N* is number of cells between the source and the sink, and *l* is length of each cell [30].

It is pertinent to mention here that an analysis based on protein lifetime measurements has shown that a temporally varying diffusivity for Bcd is somewhat better at capturing the dynamics of gradient formation for this morphogen compared to the conventional treatments that assume a constant diffusion rate [31]. However, the precise mechanism behind such temporal dependence remains unknown. The quantum walk model provides an explanation of this observation by showing that if the system can vary its diffusivity depending upon its requirement by modulating noise, then there is no single fixed value of local diffusivity for the system but rather a range of possible values. What this means, in other words, is that the Bcd protein mobility changes with time and spatial position in the embryo as is found experimentally to be the case as well [11].

As per the quantum walk analysis the diffusivity of *Bcd*, Eq. (2), is very large for the observed low noise levels in the system and the system can vary its diffusivity depending upon its requirement by modulating noise [12]. It is thus quite conceivable that a significant fraction of *Bcd* molecules can move much faster given the very low observed degradation rates, *k* ∼ 10^−4^*s*^−1^, of the *Bcd* morphogen in the system [32]. Therefore, substituting Eq. (2) in Eq. (1) one obtains

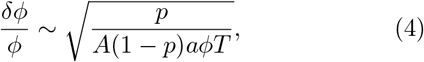

Equation (4) is our modified Berg-Purcell limit for the *Bcd* gradient. We now use it to obtain the expected *Bcd* interpretation time in order for it to be efficiently decoded by its primary target gene *hb*. For the numerical values of noise *p* comparable in magnitude to the observed *Bcd* degradation rates (*k*∼ 10^−4^*s*^−1^) in the system, and after substituting the values of concentration *ϕ* and receptor size *a* from ref. [13] into above equation

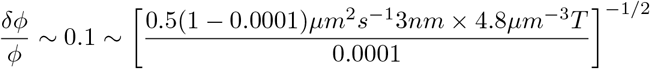

we see that the decoding of positional information from the *Bcd* morphogen gradient by its primary target gene *hb*, with the observed precision of ∼10 % takes *T*∼ 0.7 s, which is less than a second. From this one can readily calculate the on-rate of the *Bcd* morphogen for its target loci,

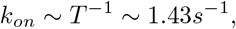

which is a very moderate on-rate. This is significant considering that lattice light sheet microscopy experiments have shown *Bcd* binding to chromatin sites in a highly transient manner, with specific binding events lasting on the order of a second, in all portions of the embryo with a rapid off rate such that its average occupancy at target loci is on-rate dependent [18].

We shall now briefly explain the observed time windows for *Bcd* interpretation. By subjecting the fly embryos to illumination at precisely defined stages, distinct time windows for the interpretation of *Bcd* have been defined through optogenetic experiments [33]. It has been demonstrated that temporal integration is indeed a necessary process in *Bcd* decoding with the integration time window covering as long as 90 minutes from nuclear cycle 10 until the end of nuclear cycle 14 [33]. Interestingly, the nuclei exposed to highest local *Bcd* concentration, those on the anterior side, integrated the signal for the longest duration while those receiving lower Bcd dosage, that is nuclei closer to the posterior side, required the inputs for comparatively shorter times in order to arrive at correct cell fates [33].

A spatio-temporally nonuniform degradation of *Bcd* could lead to a non-uniform distribution of noise in the system. Such a non-uniformity is a result of the higher concentration of *Bcd* molecules on the anterior side than that towards the posterior side of the embryo. This is despite the fact that the net concentration of *Bcd* inside the nuclei at a given position along the *AP* -axis remains constant within ∼ 10 % accuracy from nuclear cycle 10 onward [27]. The net effect of all these increased noise processes on the anterior is to reduce the diffusivity of the system, the *A*(1 − *p*)*/p* term in Eq. (4), more on the anterior side and consequently the nuclei on this side need to integrate the signal for a longer duration, higher *T* values, in order to reach the desired level (∼10 %) of precision in signal interpretation. Hence the resulting differing time windows.

The above treatment, as is clear from Eq. (4), also shows that the problems of gradient formation and interpretation are interconnected processes, at least in the case of *Bcd* − *hb* system. In this regard our treatment is a very rare instance of a long desired approach [34] that incorporates both the dynamics of morphogen gradient formation and its subsequent interpretation. Specifically, we find that the embryo does not need to rely on complex mechanisms like spatial averaging [14] and dynamically changing time windows [26] in order to generate precise boundaries of gene expression in a short span of time. On the other hand, very large values of diffusivity attained by the system due to underlying quantum effects are sufficient to achieve the desired level of precision and accuracy in the patterning process.

In conclusion we have used the recently proposed quantum walk model of *Bcd* gradient formation to explain the timely interpretation of the *Bcd* morphogen by its primary target gene *hb* with the observed precision of ∼ 10 % [13] and certain key observations of recent optogenetic experiments concerning the time windows for *Bcd* interpretation are also explained by the said model [33]. We find that the very large values of *Bcd* diffusivity are responsible for generating precise boundaries of gene expression in such a short span of time. This is in contrast to the conventional treatments that require complex mechanisms like spatial averaging and dynamically changing time windows to achieve the desired level of precision in the patterning processes. This shows that the incorporation of quantum effects into the dynamics of *Bcd* gradient formation and interpretation represents a viable step forward in exploring the biophysics of this prototypical morphogen gradient, and perhaps other morphogens like the Dpp gradient in the developing fly wing [35], in light of recent experimental observations on the system.

